# MicrobioLink: An integrated computational pipeline to infer functional effects of microbiome-host interactions

**DOI:** 10.1101/837062

**Authors:** Tahila Andrighetti, Balazs Bohar, Ney Lemke, Padhmanand Sudhakar, Tamas Korcsmaros

**Author notes:** Current address: Laboratory of Structural Bioinformatics and Computational Biology (SBCB), Institute of Informatics, Federal University of Rio Grande do Sul (UFRGS), Av. Bento Gonçalves 9500, 91501-970 - Porto Alegre, RS - Brazil. Joint corresponding authors, **Corresponding authors**, Tamas Korcsmaros, Group Leader, Earlham Institute, Norwich Research Park, NR4 7UZ, Norwich, UK, Padhmanand Sudhakar, Earlham Institute, Norwich Research Park, NR4 7UZ, Norwich, UK.

## Abstract

Microbiome-host interactions play significant roles in health and in various diseases including auto-immune disorders. Uncovering these inter-kingdom cross-talks propels our understanding of disease pathogenesis, and provides useful leads on potential therapeutic targets. Despite the biological significance of microbe-host interactions, there is a big gap in understanding the downstream effects of these interactions on host processes. Computational methods are expected to fill this gap by generating, integrating and prioritizing predictions - as experimental detection remains challenging due to feasibility issues. Here, we present MicrobioLink, a computational pipeline to integrate predicted interactions between microbial and host proteins together with host molecular networks. Using the concept of network diffusion, MicrobioLink can analyse how microbial proteins in a certain context are influencing cellular processes by modulating gene or protein expression. We demonstrated the applicability of the pipeline using a case study. We used gut metaproteomic data from Crohn’s disease patients and healthy controls to uncover the mechanisms by which the microbial proteins can modulate host genes which belong to biological processes implicated in disease pathogenesis. MicrobioLink, which is agnostic of the microbial protein sources (bacterial, viral etc), is freely available on GitHub (https://github.com/korcsmarosgroup/HMIpipeline).

## 1. Introduction

Microbiota-host interactions happen in almost every known organism, shaping their metabolism and evolution [1,2]. In many ecosystems, the microbiome plays an important role as manifested by its dynamic interactions with different hosts [3]. The community of microorganisms are almost indispensable to human life since they modulate and influence immunity and nutrient acquisition. For example, the gastrointestinal microbiome plays a crucial role in nutrient assimilation and energy yield by actively participating in metabolic pathways [4]. Dysbiosis (compositional alterations) of gut microbial communities is associated with diseases such as type 2 diabetes, obesity and inflammatory bowel diseases like Crohn’s disease [5,6]. In addition to infections caused by pathogenic microbes, exclusion of beneficial species from the community are also known to have negative impacts on the host [7]. Most of the inferences between microbial composition changes and disease phenotypes have been based on associations and correlations, with little explanations of the mechanisms driving the phenotypes.

Studying microbiome-host interactions and its influence on host biological mechanisms is important for monitoring health and disease and also for discovering/fine-tuning therapeutic interventions. Such interactions are mediated by the interplay between various molecular components expressed by the host and the microbiome. For example, bacterial molecules such as metabolites [8], proteins [9–12] and small RNAs [13], can interact with host molecules, and through host intracellular pathways and regulatory networks, modulate the expression of genes in various biological processes [14–16], thus maintaining healthy states or transition to diseased states. Besides the metabolite mediated interactions, protein-protein interactions (PPIs) are one of the most relevant types of molecular interplay between microbes and host organisms [12]. Experimental techniques to test inter-species PPIs are time consuming and have limitations imposed by cost [1,17–21]. Hence, there is a dearth of validated inter-species PPIs in publicly available databases, and these are mostly limited to pathogens. However, using predicted microbe-host PPIs, it is possible to build interaction networks to better understand the cross-talk between the microbial and host proteins, and how these interactions interfere with host metabolism and physiology [22]. Recent studies have also shown how protein-protein interactions between emerging pathogens such as COVID-19 and the host have enabled the discovery of potential drug candidates for clinical testing and validation [23].

Various resources and pipelines which generate or store microbe-host interaction predictions are currently available. They include PHISTO [24], PATRIC [25], Proteopathogen2 [26], VirBase [27] and HPIDB [28]. Most of these resources store PPIs and/or genomic analysis of virulence, but are limited to pathogens [22]. Tools such as COBRA [29], RAVEN [30], NetCooperate [31] and Kbase [32] which consider commensal microbes are based on the use of metabolomics data alone [33]. Moreover, most of the existing resources and pipelines are confined to predicting the direct molecular interactions at the microbe-host interface, and do not infer the downstream effects on functional processes and host signalling pathways.

To fill these gaps in understanding the effect of the microbiome on the host, we introduce MicrobioLink - a computational pipeline to analyse microbiome-host interactions at a cellular level using network and systems biology approaches. Using MicrobioLink, it is possible to identify potential pathways by which microbes can modulate the expression of key host molecules such as genes, proteins or microRNAs. In order to demonstrate the applicability of MicrobioLink, we performed a case study investigating how the gut microbiome potentially modulates autophagy genes in Crohn’s disease (CD). By providing custom-made lists of microbial proteins, host receptor proteins, molecular interaction networks and the target node sets, users can harness the functionalities of MicrobioLink to understand the mechanisms which mediate the influence of microbial proteins derived from the individual microbes or the microbiome.

## 2. Materials and Methods

MicrobioLink enables users to integrate host responses into the interaction networks representing the microbe-host interface, and thereby, helps users to expand the list of experimentally verifiable hypotheses. It provides the user with the option to infer indirect effects of microbial proteins on the host by prioritizing pathways and signalling chains. This prioritization is based on various criteria such as chain length, i.e, the number of steps between the host receptors modulated by the microbial protein and the host target genes/proteins, host protein localization (depending on the mode of infection) and contextual gene expression.

### 2.1. Compilation of input proteins and genes

The first step of the workflow involves the compilation of bacterial and host proteins that can potentially interact under the studied biological conditions (**Figure 1**). The bacterial proteins can be obtained from annotated or experimentally derived bacterial proteomes, or from metaproteomic read-outs (set of proteins secreted by an entire microbial community). As for the host proteins, the list is compiled based on their localization: the chosen proteins must be located in a cellular compartment which is physically accessible to the bacterial proteins. Localization based filtering can be applied also to bacterial proteins depending on the context (based on whether the microbe is extracellular or intracellular within the host). In the case of extracellular microbes, the provided host proteins must be present at the extracellular matrix or plasma membrane since these locations are more prone to interacting with extracellular, secreted or membrane bound bacterial proteins. In the case of intracellular microbes, the provided host proteins must be located inside the host cell, as in the cytoplasm for example. These host proteins provided by the user can be derived either from experimental data or publicly available datasets (including but not limited to ProteomicsDB [34], ComPPI [35], Human Protein Atlas (HPA) [36] and MatrixDB [37]). In addition, the localization of host proteins can be inferred by bioinformatic tools such as PSORTdb[38], SignalP [39] and Secretome [40], which use sequence-based features to predict the localization of the given proteins. Users can also provide their own pre-processed lists of host proteins after localization prediction.

**Figure 1:**
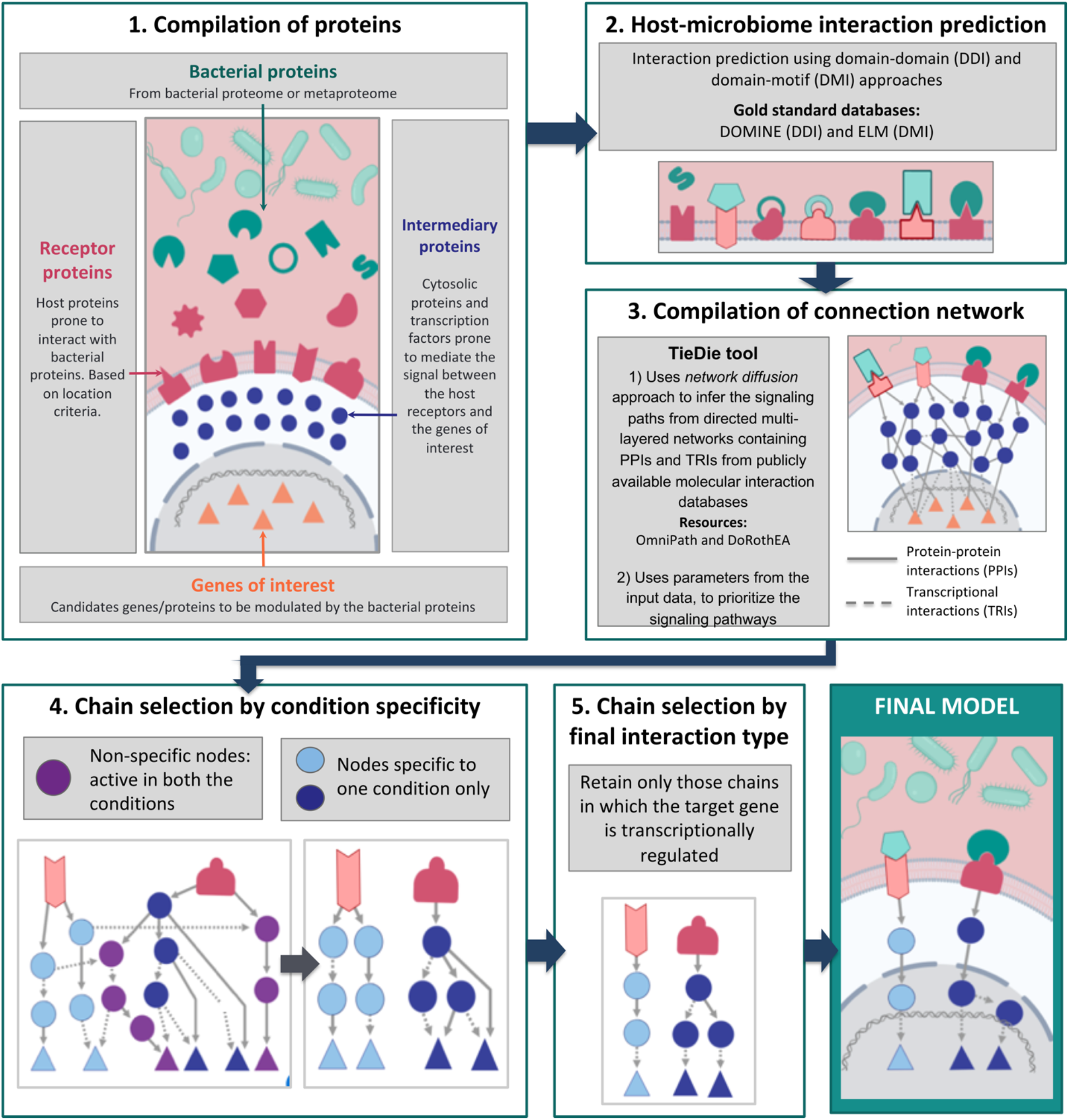
Graphical representation of the MicrobioLink workflow.

Subsequently, users can compile a list of important proteins or genes for the studied conditions or related cellular processes for use as target nodes within the host. This target list can be compiled either from *a priori* knowledge (derived from phenotypic observations) or contextual data obtained from genome-wide association studies (GWAS) which correlate genetic loci to observed phenotypes, literature search or -omic expression datasets such as gene expression (transcriptomics) or proteomics measured under the studied condition(s).

### 2.2. Bacterial-host interaction prediction

The next step in the pipeline involves the interaction prediction between the microbial and host proteins. We use two qualitative approaches, namely domain-domain and domain-motif methods [12,41,42], that use secondary structure based features to predict the interactions between microbial and host proteins. The domain-domain method is based on the assumption that microbe-host protein pairs bearing interacting domain pairs, also interact. Similarly, the domain-motif method enables the identification of potential interactions between microbial and host proteins if the microbial protein contains domains previously known to interact with eukaryotic motifs on the host proteins. Gold standard information on interacting domain pairs and interacting domain-motif pairs are retrieved from the DOMINE [43] and ELM [44] databases.

The above mentioned methods help determine which bacterial proteins, *via* their respective domains, interact with host receptor proteins. For quality control purposes, the interactions are then filtered to sterically possible sequence features [45,46]. This is performed by excluding interactions involving motifs outside disordered regions (using IUPRED [47]) or within globular domains based on information from PFAM [48] and InterPro [49]. Previous studies have successfully used this structure-based approach to predict PPIs between microbial and host proteins, and validated the PPIs experimentally [9,10,12].

### 2.3. Network compilation and path tracing using diffusion

In this step, we use multi-layered network resources and network diffusion tools to trace the effect of the interactions between microbial and host proteins on other host processes further downstream. For this purpose, the TieDIE [50] tool, built into the pipeline, is used to infer the signaling paths that connect the host receptors with the host target genes. TieDIE uses the network diffusion approach to infer the signaling paths from directed multi-layered networks containing PPIs and transcriptional regulatory interactions (TRIs) from publicly available molecular interaction databases such as OmniPath [51] and DoRothEA [52] respectively. Users can also select from a vast number of other molecular interaction resources which are available in the public domain. The use of high-quality curated PPI resources such as OmniPath enables the analysis of post-translational modifications as many original sources in OmniPath contain such information. Furthermore, integration of proteomics and/or phosphoproteomics data to be analyzed in the MicrobioLink pipeline can improve the computational predictions by imparting activity information into signalling proteins and prioritizing the predictions (i.e activated signalling paths) for experimental validation.

Importantly, the network diffusion approach helps in avoiding prioritization biases stemming from network topological properties such as degree (i.e the number of interacting first neighbours of a particular gene or protein for example). This circumvents the positively biased inclusion of hub proteins into signalling paths irrespective of their actual relevance in the studied condition [50]. TieDIE also incorporates various parameters along with the input data. For example, the magnitude and direction of differential expression of the target genes/proteins or the number of microbial proteins predicted to bind to the host receptors can be specified into the input dataset. This information is then used to prioritize the signaling pathways connecting the receptors to the target genes/proteins.

After network compilation, depending on the user’s discretion, a chain selection step can be performed to select the most relevant chains for the target modulation. This can be done by filtering the network to keep only signalling chains stimulated by bacterial proteins detected in any single condition, especially in cases in which the users are comparing different conditions. Chain selection can also be specified by the user depending on the dataset provided. For example, in our use case, the host expression dataset is determined from transcriptomic datasets. Hence, we confined the interactions immediately upstream to the target genes to transcriptional regulatory connections, in order to capture the regulatory biology underlying the modulation of the target genes’ expression.

Finally, the obtained model can be visualized as a multi-layered network, starting with the bacterial protein-host receptor interactions, and ultimately reaching the selected host target genes through PPIs and TRIs. A summary of the MicrobioLink pipeline is presented in **Figure 1**.

## 3. Results

### 3.1. Case study

To demonstrate the applicability of the MicrobioLink pipeline, we performed a case study showing how different bacterial proteins modulate molecular processes important for the development or progression of Crohn’s disease (CD). CD is a chronic intestinal inflammatory disease (IBD) associated with a multitude of factors including microbial dysbiosis and dysregulation of autophagy due to mutational effects [53–55] [56,57]. Using MicrobioLink, we identified pathways by which autophagy could be indirectly modulated by microbial proteins, thereby implying that host autophagy can be potentially modulated by microbial influence independent of mutations. This also highlighted the potential role of the microbial proteins in disease development and/or progression.

To perform this study, we first obtained the microbial proteins present only in healthy individuals or in CD patients from a Swedish twin study [58]. In total, 336 microbial proteins from healthy individuals and 226 proteins from CD patients were obtained (**Supplementary table 1**). Then, we compiled human proteins located in the extracellular matrix and cellular membrane using matrixDB, resulting in a total of 8008 proteins (**Supplementary table 2**).

After compiling the protein lists, we performed interaction predictions between microbial and human proteins, resulting in 8478 predicted interactions involving 140 microbial proteins and 2998 host receptor proteins (**Supplementary tables 3-4**). The predicted interactions were refined by passing them through a disordered region based quality control step to eliminate sterically improbable interactions. In parallel, as potential target genes affected by the microbial proteins, we focused on autophagy genes given that autophagy is known to be a dysregulated process in CD [53–55,59,60]. In total we selected 38 autophagy genes, encoding the core components of the autophagy machinery [61]. To detect the most relevant target autophagy genes for CD, we used three different transcriptomic datasets (GSE9686 [62], GSE36807 [63] and GSE75214 [64] from GEO [65]), that contained the gene expression profiles of CD patients and healthy individuals, to select the most differentially expressed core autophagy genes. From the above mentioned profiles, we selected autophagy genes which were modulated with coherent trends in at least two datasets. Based on this criteria, we obtained five target autophagy genes namely WIPI1, MAP1LC3A, MAP1LC3B, ATG7 and ATG4D (**Supplementary table 5**).

Subsequently, contextual signaling networks that potentially mediate the signal transduction from the host receptors to the target autophagy genes were compiled using the network diffusion model inferred by TieDie (**Figure 2A**). Based on these results, we were able to infer that two of the five target autophagy genes could be potentially modulated by the microbial proteins. We also observed a group of host proteins that are connected to microbial proteins occurring in both CD patients and healthy subjects. To retain specificity, we excluded the chains initiated from microbial proteins enriched in both conditions. As a result, we obtained a network with clearly separated signalling paths exclusive to the disease and healthy contexts as shown in **Figure 2B**. In a final filtering step, we retained only those chains where the last interaction is a TRI to capture the transcriptional regulatory effect of the expression of the target genes. Post this final filtering step, we were able to obtain a final model, as depicted in **Figure 2C**.

**Figure 2:**
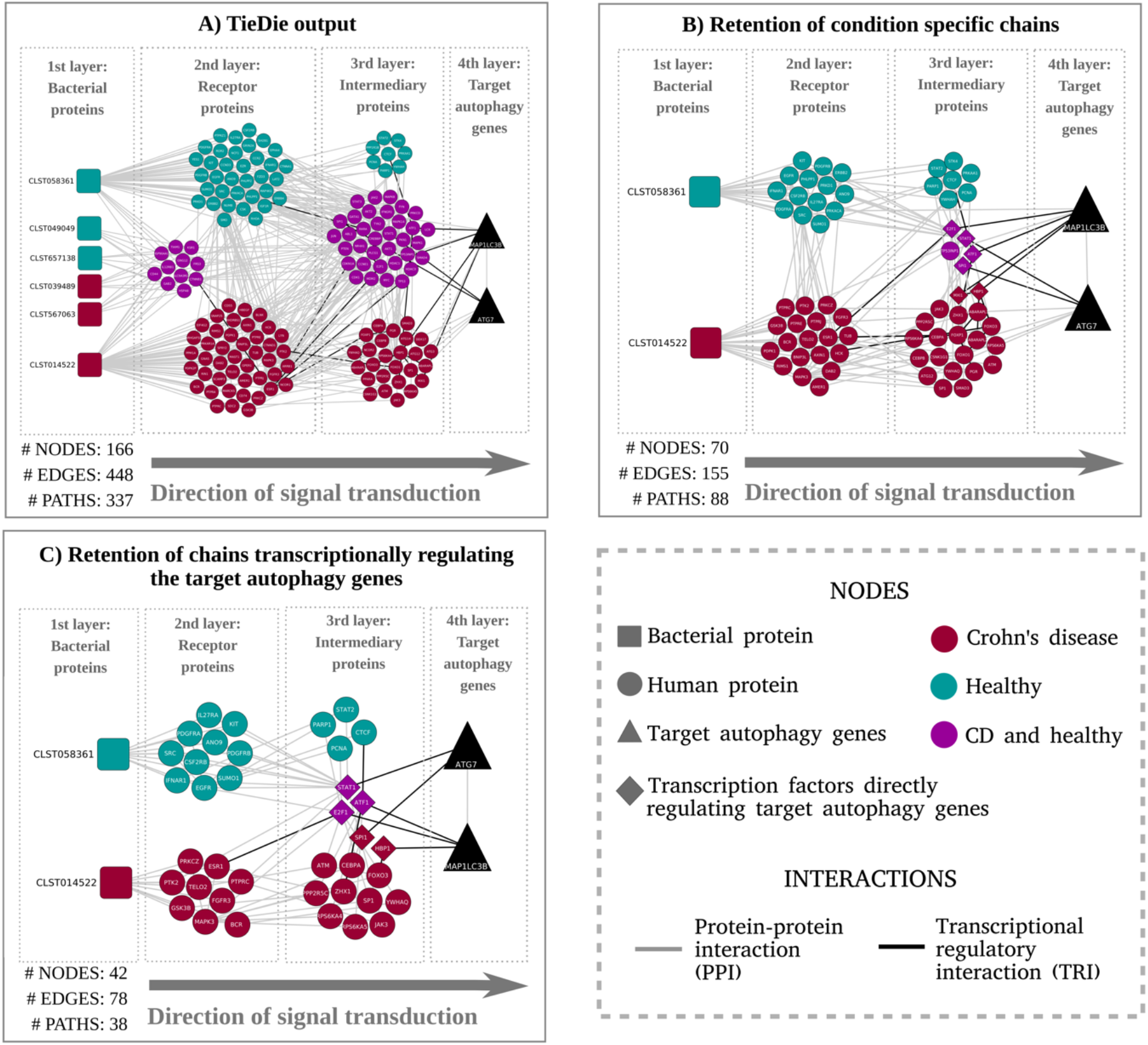
**(A).** Graphical representation of the signaling paths between curated receptor proteins (host proteins predicted to be modulated by microbial proteins) and target autophagy genes. The network compilation was performed by tracing the signaling chains from the human receptors (predicted to interact with the bacterial proteins) to the autophagy genes using the TieDIE tool which adopts a network diffusion [51]. For brevity, only the results corresponding to the domain-motif interaction analysis are discussed in the case study. **(B).** Network representing the signalling chains after exclusion of proteins connected with bacterial proteins detected in both CD and healthy conditions. Proteins present in both conditions that were retained were those directly regulating the target autophagy genes. **(C).** Network obtained by retaining only chains with transcriptional regulatory interactions between the intermediary protein (3rd layer) and the target autophagy genes (4th layer). The immediate upstream proteins from the autophagy target genes were confined to transcription factors modulating the target autophagy genes via a transcriptional regulatory interaction.

To analyse the functional significance of the final model, we did a Gene Ontology (GO) enrichment analysis of the human proteins to identify feature sets such as biological processes *via* which the target autophagy genes are potentially modulated by the microbial proteins. Over-represented feature sets (**Figure 3, Supplementary table 7**) included apoptosis, which is known to be upregulated in intestinal epithelial cells (IECs) in IBD patients leading to increased epithelial barrier permeability [66]. Apoptosis has a peculiar interplay with autophagy, in that it is usually up-regulated when autophagy is deactivated and *vice versa* [67]. In our final model, we also discovered several signaling chains among host networks specific to the healthy condition. These signalling chains included proteins such as SUMO1, PARP1 and E2F1. SUMO1, known to induce autophagy levels [68], also activates PARP1 and subsequently E2F1, proteins which are both known to stimulate autophagy and inhibit apoptosis [69,70]. This allowed us to propose the hypothesis that the metaproteome-derived bacterial protein CLST058361 which was expressed in healthy subjects and not in CD patients, could be a key protein involved in stimulating autophagy and decreasing apoptosis. Interestingly, CLST058361 belongs to a family of trypsin-like serine proteases whose expression level changes are associated with IBD [71] [72] [73],

**Figure 3.**
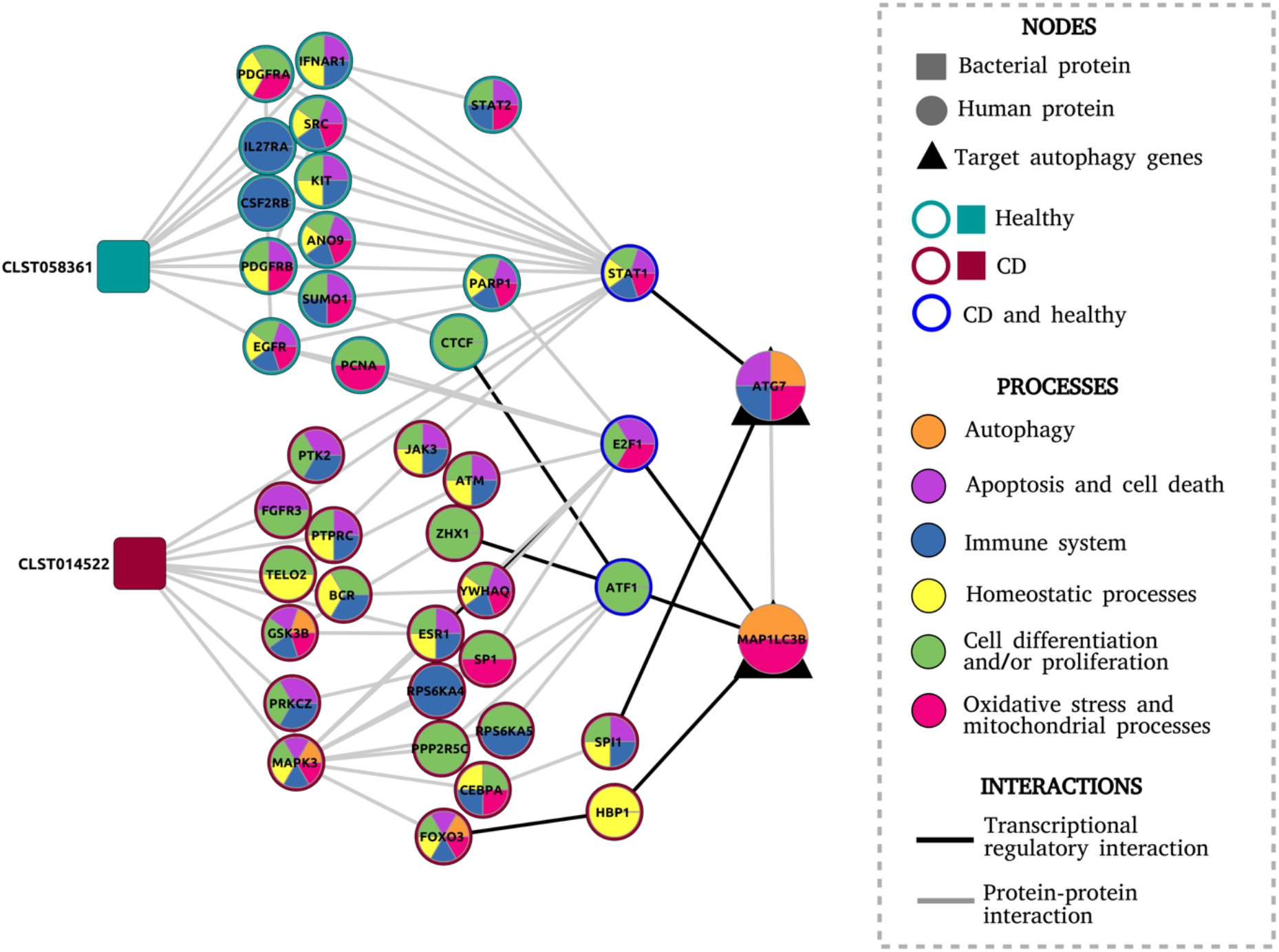
Final network model consisting of the proteins and the biological processes in which they are involved, inferred from the GO enrichment test.

Meanwhile, a signalling chain specific to the CD condition included MAPK3 and FOXO3, which are key proteins for apoptosis activation and apoptosis-autophagy interplay [67,74]. This pathway was modulated by the bacterial protein labeled as CLST14522 (and containing a FOG:_FHA_domain) and uniquely found in CD patients. Other biological processes over-represented in CD networks included those related to cellular differentiation and proliferation, response and generation of oxidative stress, mitochondria related processes, homeostasis and immune system regulation. More comprehensive studies are warranted to ascertain the role of these processes in how the bacterial proteins modulate autophagy in CD (**Figure 3**).

In conclusion, using MicrobioLink, we obtained a mechanistic model showing how bacterial proteins can modulate the expression of autophagy genes in the context of CD. Previous research has shown that host autophagy, while being implicated in the pathogenesis of CD, was primarily dysregulated by defects caused by genetic mutations [53–55] [56,57]. MicrobioLink reveals the possibility that autophagy can be modulated by the microbial community through signal transduction propagated *via* host molecular networks. Although experimental validation is required to ascertain this putative mechanism, we demonstrate the applicability of MicrobioLink to integrate heterogeneous datasets and generate testable hypotheses about microbe-host interactions.

## 4. Discussion

In this paper, we presented MicrobioLink - a pipeline for the analysis of the functional effects of the microbiome on host cellular processes. MicrobioLink integrates microbiome-host protein-protein interaction predictions with network diffusion to infer signaling networks which capture the systemic effects of microbial proteins on host processes.

Microbiome-host interactions have considerable impact on host signaling and understanding these interactions are crucial to our advances in studying disease pathogenesis and discovering drug targets. Various computational tools and approaches have enabled and furthered our understanding of microbiome-host interactions. For example, NetCooperate [31] uses metabolic network modelling to infer the nutritional interdependencies and co-feeding potential between any two microbial species or between microbial and host species. Zanzoni et al [75] harnessed the power of microbe-host protein-protein interactions to infer the perturbations induced on the host by the pathogen *Fusobacterium nucleatum*. However, researchers aiming to study the functional effects of microbiome-host interactions face technological and methodological boundaries. First, the high demand of time and cost of experimental studies makes it infeasible to test such a high number of potential microbiome-host interactions [19]. MicrobioLink is an effective alternative to study the molecular mechanisms by which microorganisms interact with the host cells and how these interactions affect the host cellular mechanisms. It can be applied either as a tool for prioritizing the most relevant non-canonical *de novo* pathways or for establishing hypotheses about how microorganisms can modulate particular cellular processes. Other limitations that MicrobioLink helps overcome include the possibility for the user to customize the pipeline depending on his/her organism or community of interest as long as the focus of the microbial-host molecular interface are protein-protein interactions. Due to the mechanism driven approach of MicrobioLink, it provides the users with experimentally testable hypotheses.

Several existing computational methods and tools to study microbe-host interactions are currently available although they are limited to specific organisms (with a focus on pathogens) and specific types of interactions. Therefore, to perform an analysis where the microorganism-host interaction is unknown and to study its influence on host processes, the researcher would have to combine different and incompatible tools. In **Table 1**, we benchmarked some of the key resources, and compared them with the functions of MicrobioLink. With MicrobioLink, it is possible to perform functional microbiome-host analysis without previous knowledge about its interactions for any microorganism and host species as long as the corresponding annotated proteomes/genomes are available. The user can provide a protein dataset as input from a single species (proteome) or from an entire microbial community (metaproteome). These microorganisms can be bacteria, archaea, viruses or fungi, which makes MicrobioLink applicable for any type of microorganism. Subsequently, the user can select a dataset that represents the resulting host phenotype. It is possible to choose any dataset adequate for the study, such as differentially expressed genes from transcriptomic measurements or protein expression from proteomics. The host data can also correspond to any host, and hence MicrobioLink can be applied for studying the influence of the microbiome on plants, mammals or even fungi.

**Table 1:** Summary of selected tools and resources used in microbe-host interaction research.

| Resource/<br>tool | Standalone<br>version ? | Description | Can user -<br>provided<br>datasets be<br>handled ? | Non-<br>pathogenic<br>species<br>included /<br>handling ? | Protein -protein<br>interactions ? | Inferring<br>downstream<br>effects ? | Microorgani<br>sms<br>supported | Host<br>organisms<br>supported |
| --- | --- | --- | --- | --- | --- | --- | --- | --- |
| PHISTO<br>[24] | online | Web-tool for<br>mining and<br>retrieving<br>host-<br>pathogen<br>interactions | no | no | yes | no | Viral,<br>bacterial,<br>fungal and<br>protozoan<br>pathogens | Human |
| PATRIC<br>[25] | online | Genome<br>focussed<br>infectious<br>disease<br>research<br>database | yes | no | yes | no | Bacterial<br>pathogens | Actinopter<br>gii,<br>Arachnida<br>,<br>Chromado<br>rea,<br>Insecta,<br>Mammalia |
| Proteopat<br>hogen2<br>[26] | online | Database and web application to store and display fungal pathogen proteomics data. | no | no | no | no | Fungal pathogens | Mammalian species |
| VirBase<br>[27] | online | Database of virus-host ncRNA-associated interactions and interaction networks during viral infections. | no | yes | no | no | Virus | Vertebrates, plants and arthropods |
| NetCoperate<br>[31] | python module | Web-based tool and software package for determining host-microbe and microbe-microbe cooperative potential from metabolic networks. | yes | yes | no | yes | Any microorganism | Any host species |
| Kbase<br>[32] | Online, python and java | Software and data platform that enables data sharing, integration, and analysis of microbes, plants, and their communities by creating workflows consisting of a series of analysis tool runs<br>and code<br>blocks | yes | yes | no | yes | Any microorganism | Any host |
| M<sup>2</sup>IA<br>[80] | web-based<br>server | Statistical<br>analysis<br>methods for<br>microbiome<br>and<br>metabolome<br>data<br>int<br>egration,<br>including<br>correlation<br>an<br>alysis and<br>functional<br>network<br>analysis | yes | yes | no | yes | Any<br>microorgani<br>sm | Any host<br>species |
| COMETS<br>[81] | Matlab and a<br>python<br>toolbox | Modelling<br>framework<br>that<br>integrates<br>dynamic<br>flux balance<br>analysis<br>with<br>diffusion to<br>communitie<br>s | yes | yes | no | yes | Any<br>microorgani<br>sm | Any host<br>species |
| Microbiol<br>ink (this<br>paper) | Python,<br>Docker | Integrated<br>evaluation<br>of microbe-host<br>interaction<br>networks | yes | yes | yes | yes | Any<br>microorgani<br>sm | Any host<br>species |

For reproducibility and interoperability [76], we have implemented the pipeline within the framework of the Docker system. Docker functions as an integrated platform with all the needed dependencies and software packages pre-installed to execute the scripts within the pipeline. Users can thereby quickly test different components of the pipeline within the docker container without having to search and install the required dependencies. The datasets corresponding to the use-case have also been provided within the container so that the users can execute the pipeline readily and cross-check the results.

The pipeline can be applied to different diseases modulated by the microbiome. In the case study, we generated a model explaining how the gut microbiome from CD patients is potentially modulating autophagy, a cellular process known to be altered and dysregulated in CD patients. Besides, the tool can be extended to diverse diseases that are shown to be influenced by the microbiome such as diabetes, depression and cardiovascular disorders [77]. Moreover, beneficial effects can also be studied with MicrobioLink. In humans, it can be applied to demonstrate how probiotic species maintain health and homeostasis. In livestock and poultry, it can be used to study how the microbiome can influence productivity [78]. In plants, MicrobioLink can be used to analyse how the commensal microbiome in the soil influences plant health and disease [79]. Eventhough experimental validation of the inferences from the pipeline is out of the scope of this work, we previously demonstrated [12] the validity of the computational inference of microbe-host protein protein interactions and their downstream effects by *in vitro* verification. As a next step towards improving the pipeline, we are planning to establish use cases in different domains of biological research (including aiding generation of hypotheses for experiments) wherein the microbiome plays a crucial role in the phenotypes of different hosts such as plants as well as mammalian host species. In particular, MicrobioLink will be very useful in studying the role of the microbiome in autoimmune disorders, such as rheumatoid arthritis, vasculitis, multiple sclerosis and systemic lupus erythematosus. With MicrobioLink, we could infer the microbiome-mediated mechanisms in such disorders and thereby point out key microbial inferences, cellular pathways transmitting normal microbial signals affected in these disorders, and potential targets for further therapeutic interventions.

## 5. Conclusion

For many conditions and organisms, MicrobioLink presents novel additional functionalities which makes it possible not only to predict microbiome-host interactions but also to infer mechanisms driving functional effects further downstream. MicrobioLink provides an integrated approach by embedding the host proteins modulated by the user-provided microbial proteins within molecular interaction networks and relevant -omic datasets customized for the studied conditions and organisms. Using MicrobioLink, it is possible to evaluate how an entire microbial community or even a single microorganism, either a commensal or pathogen, can interfere with host processes *via* protein mediated signal transduction.

## Data and Software Availability

Code for the MicrobioLink pipeline is available at https://github.com/korcsmarosgroup/HMIpipeline. The code is implemented in Python and dockerized in order to ensure easy testing, circumvent dependency issues and achieve reproducibility. The implementation is provided as two versions - firstly as a sequential step-by-step workflow and secondly as a one-click version. Both versions have been supplemented with corresponding documentation including the dependencies and instructions for execution.

## Author contributions

TK and PS designed the methodology; PS and TA developed the codes; TA and PS performed the case study; BB performed the dockerization. TA, TK, NL and PS wrote the manuscript; all authors read and contributed to the manuscript.

## Supplementary tables

**Supplementary table 1**. List of microbial proteins derived from the Swedish twin cohort study [58].

**Supplementary table 2**. Localization filtered human receptor proteins retrieved from MatrixDB [37].

**Supplementary table 3**. Domain-domain based protein-protein interaction predictions between microbial and human receptor proteins.

**Supplementary table 4**. Domain-motif based protein-protein interaction predictions between microbial and human receptor proteins.

**Supplementary table 5.** Gene expression measurements, as represented by the logFCs, of the core autophagy genes from three different GEO datasets.

**Supplementary table 6**. Signaling network obtained by TieDIE [50], connecting the host receptor proteins (predicted to bind to the microbial proteins) and the target autophagy genes.

**Supplementary table 7**. GO enrichment results of the human proteins from the final network model. The 1st sheet contains all enriched gene ontology terms (biological process) in the final network. The 2nd sheet contains selected processes related to CD pathogenesis. The 3rd sheet contains the assignment of proteins to the selected biological process terms enriched in the network.

